# Counterconditioning following memory retrieval diminishes reinstatement of appetitive memories in humans

**DOI:** 10.1101/376392

**Authors:** Rani Gera, Segev Barak, Tom Schonberg

**Author notes:** Corresponding Authors: Tom Schonberg, Faculty of Life Sciences, Department of Neurobiology and The Sagol School of Neuroscience, Tel Aviv University, Tel Aviv 69978, Israel; Segev Barak, School of Psychological Sciences and The Sagol School of Neuroscience, Tel Aviv University, Tel Aviv 69978, Israel. **Competing Interests:** The authors declare no competing interests.

## Abstract

Appetitive memories play a crucial role in learning and behavior but under certain circumstances become maladaptive and play a vital part in addiction and other psychopathologies. In recent years, scientific research demonstrated the ability of memories to be modified following their reactivation through memory retrieval, in a process termed memory reconsolidation. Several non-pharmacological behavioral manipulations yielded mixed results in their capacity to alter maladaptive memories in humans. Here, aiming to translate promising findings observed in rodents to humans, we constructed a novel three-day procedure to test the efficacy of aversive counterconditioning to alter appetitive memories when given following memory retrieval. On Day1 we used appetitive conditioning to form appetitive memories. On Day2, we retrieved these appetitive memories for half of the participants (Retrieval group). Subsequently, all participants underwent counterconditioning. On Day3, we tried to reinstate Day1 appetitive memories. We observed a significant reduction in the reinstatement of the original appetitive memory when counterconditioning was induced following memory retrieval. We provide here a novel human paradigm that models several memory processes, and demonstrate memory attenuation when counterconditioned after its retrieval. This paradigm can be used to study complex appetitive memory dynamics, e.g., memory reconsolidation, and their underlying brain mechanisms.

## Introduction

Appetitive associative memories play a crucial role in motivation, learning, behavior and decision-making ^1^. Over the last two decades, it has been demonstrated that associative memories can be modified when manipulated during the process of ‘memory reconsolidation’ (for review see Lee et al.^2^). In this process, reactivation of consolidated memories via their retrieval initiates a temporary labile state that lasts a few hours ("reconsolidation window"), during which memories are prone for modification before their re-stabilization ^3,4^.

Interference with the reconsolidation process can potentially disrupt or update the original memory, consequently preventing its expression, and thus holds significant clinical implications for disorders implicating associative memories such as addiction^5^. Despite substantial clinical potential, only a few studies targeted the reconsolidation of appetitive memories in humans. Several studies employed pharmacological interventions during reconsolidation of appetitive memories ^6–8^, yet a pharmacological approach may be limited due to its potential side effects.

Previous studies in rodents, exemplified the ability of behavioral manipulations to affect the reconsolidation of associative memories ^9–11^ and a few studies showed their capacity to affect the reconsolidation of human fear memory ^12,13^. A handful of studies applied behavioral manipulations during reconsolidation of appetitive memories in humans ^14–16^, yet these studies were conducted in drug addicts or heavy alcohol abusers, where the target memory pre-existed prior to the study. However, a basic-research paradigm of these processes in healthy human individuals has yet to be developed.

Here, we aimed to develop such a new human paradigm, which disrupts the reconsolidation of appetitive memories in healthy participants. A controlled laboratory task, in which the target memory is formed under fully controlled settings, is vital for understanding the fundamental mechanisms of memory dynamics. Moreover, establishing such a paradigm is essential to characterize and tune the manipulations on memory dynamics, in order to develop clinical treatments. Recently, Goltseker et al.^17^ developed a novel behavioral paradigm in mice and demonstrated that administering aversive counterconditioning training following the retrieval of cocaine-associated memories abolished the reinstatement of cocaine seeking in mice. In the present study, we aimed to translate this novel retrieval-counterconditioning animal model to a new paradigm that alters appetitive memories in healthy humans.

We designed a behavioral procedure consisting of three stages, during which appetitive memories were formed in the laboratory, then counterconditioned and finally reinstated. Aiming to disrupt the reconsolidation of these memories by behavioral interference, we reactivated them prior to their counterconditioning for one group but not the other, using a brief memory retrieval. We hypothesized that such treatment will substantially reduce the reinstatement of these memories only in the retrieval group.

## Materials and Methods

For details on the apparatus, an algorithm produced for stimuli assignment, a sequence assignment task implemented in the manipulation phases, features used to improve task realism and a task used to separate retrieval and counterconditioning see Supplementary.

### Participants

Ninety-six (63 females) healthy participants participated in the experiment. We aimed to obtain n=25 valid participants that demonstrated adequate learning based on certain criteria (see Experimental Design and Statistical Analysis below for details) in each of the 2 groups: Retrieval and No-Retrieval. The 50 valid participants (age range 18-39, mean=24.82, sd=4.52) included 35 females. Informed consent was obtained from all participants prior to participation in the experiment in return for monetary compensation (40 NIS =~ $12 USD per hour). Participants were informed that apart from the hourly fee, they may gain or lose additional monetary bonus. They were guaranteed that even if their losses would exceed their gains, calculated from the beginning of the experiment, they would still receive as a minimum the promised hourly rate. Therefore, their total compensation in each point could not be less than the hourly accumulated payment. All experimental protocols were approved by the ethics committee of Tel Aviv University. All methods were carried out in accordance with the relevant guidelines and regulations. Four participants were excluded due to software failures, and eight due to incompletion of the experiment. Thus, a total of n=84 were valid participants out of which n=50 (n=25 in each group) qualified our experimental exclusion criteria.

### Procedure

The experiment was programmed and run in MATLAB R2014b (The MathWorks, Natick, Massachusetts, USA), using Psychtoolbox3 ^18,19^, and consisted of three stages (Conditioning, Counterconditioning, Reinstatement), across three consecutive days. Participants were randomly assigned to Retrieval and No-Retrieval groups. The overall workflow of experimental procedures is illustrated in Fig. 1A. Briefly, on Day1 we first measured participants’ baseline liking toward different stimuli and assembled the CSs accordingly. Subsequently, participants underwent appetitive conditioning. On Day2, after retrieving the memory only for the Retrieval group, participants from both groups underwent counterconditioning. On Day3 we aimed to reinstate the memory that was formed through conditioning. Preferences, as well as liking ratings, were measured following the different stages of the task and probed changes in preference and liking toward the CSs. We used computerized simulated houses as the CSs, and monetary gain and loss as appetitive and aversive UCSs, respectively.

**Figure 1.**
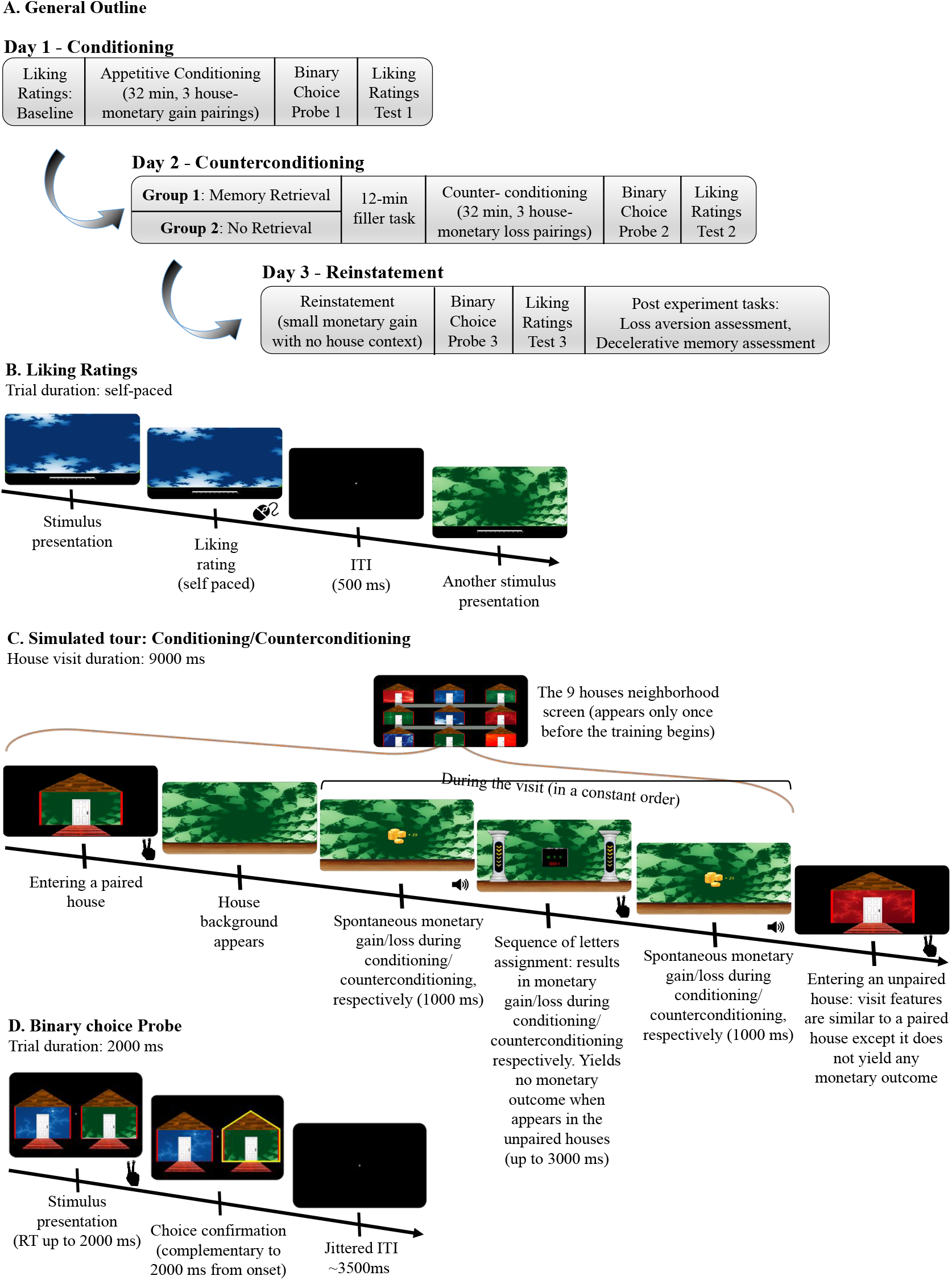
Experimental procedure outline and design. **A.** Participants attended 3 experimental stages on 3 consecutive days. **B.** Participants performed a baseline fractal liking ratings procedure. The same procedure also followed the conditioning, counterconditioning and reinstatement stages. **C.** Appetitive conditioning to simulated houses (CSs) was applied through a computerized simulated tour. In each of 20 blocks, participants visited each of the 9 houses in the “neighborhood” once in a randomized order. The houses were divided into 3 equal-sized sets, distinguished by color. One set (the green houses in this illustration) was paired with monetary gain through two Pavlovian rewards and a reinforced instrumental task occurring between them. This task was also used for the aversive counterconditioning on Day 2, in which the conditioned stimuli were paired with monetary loss. A short version of this procedure was also used for both the memory retrieval, where participants visited the paired houses and neither gained nor lost money; during reinstatement, participants visited houses with no background (stimuli), during which they received a monetary gain. **D.** In the probe parts participants had to choose which houses they preferred to enter. The images in this figure were created by the first author (RG).

## DAY 1

### Baseline fractals liking ratings

Upon arrival, participants rated their liking towards eighteen fractal art images (see https://osf.io/48uca/ for a sources list) on a visual analogue scale (VAS) ranging from 0 to 10 (Liking rating baseline) (see Fig. 1B). The fractal collection was assembled of three color sets (red, green and blue), each consist of six fractals. Fractal presentation order was randomized and the task was self-paced.

To avoid a baseline color preference effect in subsequent tasks, we implemented a designated algorithm (see Fig. 2 and Supplementary information), determining, based on participants’ subjective ratings, three fractals of each color to be used as the nine CSs (6 CS- and 3 CS+). The use of color sets was designed to allow participants to verbalize a rule, and thus increase contingency awareness, which has been shown to promote the occurrence and magnitude of conditioning (for example, see refs ^20–22^).

**Figure 2.**
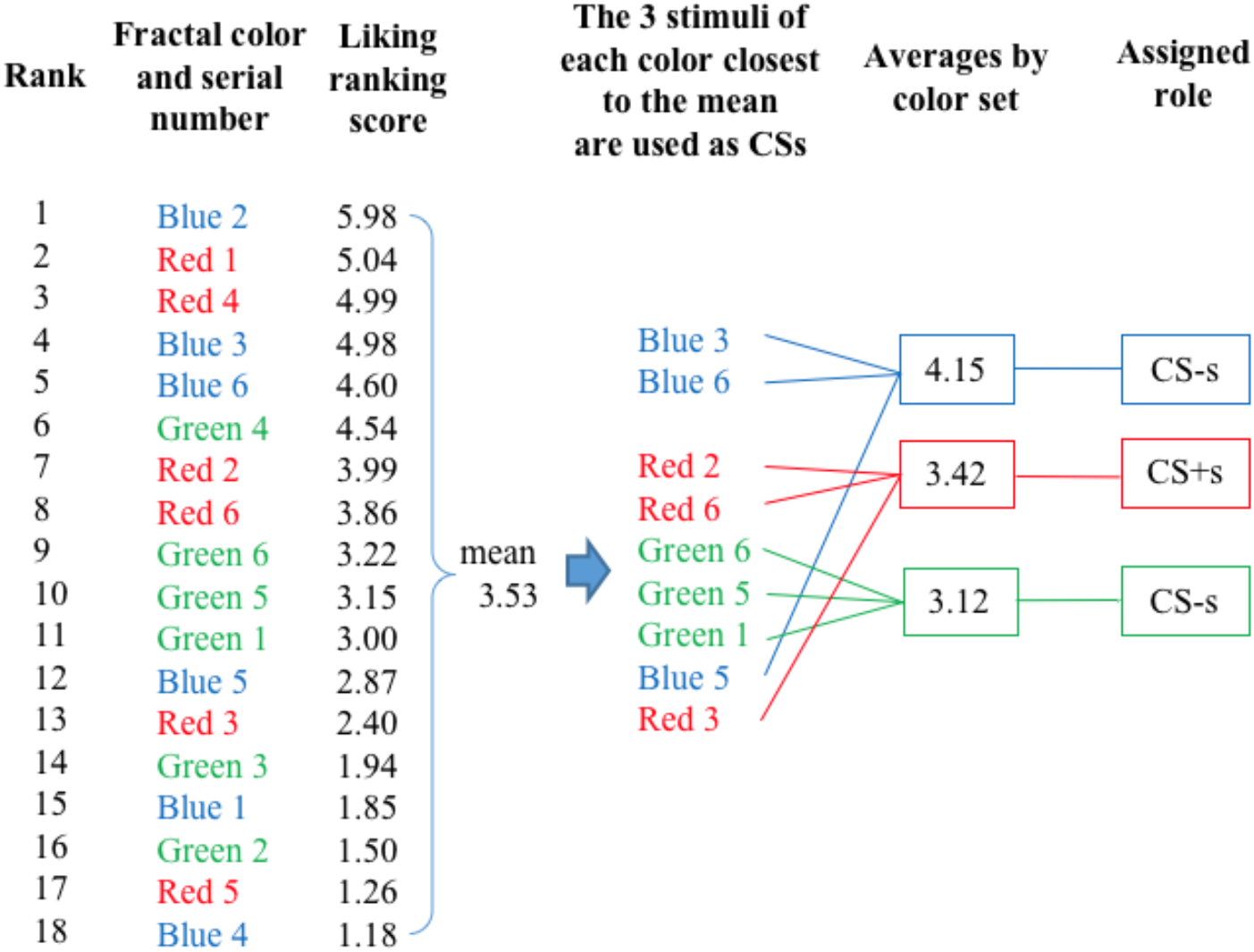
Scheme of conditioning and counterconditioning stimuli assignment. The scheme is demonstrated through one participant’s ranking scores. Stimuli were ranked and sorted according to individual ratings in the Baseline fractals liking ratings. Three stimuli of each color (red, green and blue) that are the closest to the overall ranking score mean were used together as the 9 CSs. Then, the average score for each CSs color was calculated within its group. The 6 stimuli that belonged to the colors with the higher and lower averages were used as CS-s and the 3 stimuli of the color with the intermediate average served as the CS+s. CS+ = paired stimulus, CS- = unpaired stimulus, CSs refers to both CS+s and CS-s.

### Conditioning

Participants were instructed that they will participate in a computerized simulated tour, during which they will enter different houses (see Fig. 1C). A total of nine houses were designed with an identical generic wooden floor and were distinguished only by their background wallpaper. Each of the nine fractals (CSs) served as the background of one house. Participants were instructed that in some of the houses they will receive money whereas in other houses, money will be taken away from them.

The task consisted of 20 blocks. On each block, participants visited each of the nine houses once in a randomized order, spending 9 seconds inside a house on each visit. A third of all houses (=three houses), members of the same color set, were assigned as CS+ and paired with monetary gain. On each visit of a paired house, a monetary reward (of 0.18 NIS, 5 cents on average) was delivered twice, indicated by a short animation of coins and the sum gained, popping up, accompanied by a winning casino-like sound (See Fig. 1C). In the other houses (CS-), no monetary gain nor loss was delivered.

In addition, we implemented an instrumental gaming-like sequence assignment, scheduled to occur once in each house visit (see Fig. 1C and Supplementary Information). In the paired houses (CS+) it appeared between the two reward presentations. During the conditioning stage, success in an assignment in the paired houses (CS+) yielded monetary gain and in the unpaired houses (CS-) yielded no monetary outcome. These sequence-pressing assignments were designed to increase participants’ engagement and enhance conditioning efficacy with an additional appetitive component that is paired with the paired houses (CS+). The total duration of the conditioning stage was ~32 minutes including transition between houses (excluding ~4 minutes instructions and task demonstration).

To facilitate learning we used a constant presentation order within each 9-second visit of each paired house: spontaneous reward (Pavlovian component), sequence assignment (instrumental component), and another spontaneous reward (Pavlovian component). To improve the realistic narrative of the simulated tour we implemented several designated features (see Supplementary Information for a detailed description). For example, the transition between houses was indicated by the presentation of the next house to which the participant was about to enter, simulated from the outside, with a path leading to it and a door (Fig. 1C). To open the door participants were required to press a designated keyboard button.

Participants were not informed of the true goal of the task. Apart from encouraging them to respond as fast as they could in the *sequences assignments*, we did not explicitly direct them to notice, identify or act in any particular manner.

### Binary choice probe 1

After conditioning, participants performed a binary choice probe (preference test), to measure preferences toward the paired houses (CS+) following conditioning. Participants were presented with two houses on each trial and were given 2 seconds to choose the house they preferred to enter (see Fig. 1D). They were told that on the last day of the experiment, one of the trials will be randomly chosen and that they will enter the house they chose on that particular trial and will get to spend a few minutes inside that house. We encouraged participants to treat every pair as if it was the one that will be chosen at the end of the experiment and thus act accordingly. The probe phase included two blocks. On each block, each one of the three CS+ was compared with all six CS-, constituting eighteen comparisons (3*6). To avoid a mere-exposure effect ^23^ additional nine comparisons (3*3), each consisting of two different CS-, one from each of the two color-sets assigned as the CS-, were presented to participants on each block. Thus, each CS appeared the exact same number of times. Trial order was randomized within each block.

In order to minimize the influence of short-term memory recruited during Conditioning on choices in this task, we included a faces liking ratings task between Conditioning and Binary choice probe. The structure of the task was similar to the Fractals liking ratings, except that the participants were asked to rank 60 unfamiliar faces ^24^.

### Fractals liking ratings test 1

This test was identical to the Baseline fractals liking ratings and was used to assess changes in liking toward the stimuli that assembled the paired houses compared to the stimuli that assembled the unpaired houses following conditioning.

## DAY 2

### Memory Retrieval

Upon arrival, the Retrieval group participants performed a short simulated tour in the same neighborhood they toured on Day 1. The task was consisted of two visits to each of the three paired houses (CS+). The visits were similar to Day 1, except that (in contrast to Day 1) no money was earned during these visits, thus producing a negative prediction error. Prediction error was suggested to be vital to induce memory reconsolidation ^25–27^, therefore the instructions, as well as the presentation of the neighborhood screen, were identical to the respective components of the Day 1 Conditioning stage. This was meant to increase expectancy violation by inducing participants’ expectations for an equivalent task as in Day 1. The total duration of the retrieval task was ~1.5 minutes.

Following the completion of the Memory retrieval task by the Retrieval group, participants watched a short nature video (~12 minutes including instructions) to provide a temporal separation between memory retrieval and subsequent counterconditioning. This was done to induce destabilization of the existing memory ^5^. While watching the video, participants were asked to answer several questions regarding its content (see Supplementary Information). Participants in the No-Retrieval group performed the movie part upon arrival without the memory retrieval phase. They performed this part at a nearby building with a different experimenter to avoid unintentional memory reactivation via physical context ^28^. Upon completion of the movie, they were instructed to go to the original testing room in the laboratory to continue the experiment.

### Counterconditioning

The counterconditioning stage on Day 2 was similar to the Conditioning stage in Day 1, except that the previously paired houses (CS+) were paired with monetary loss rather than gain, with equivalent amount and contingency as in Day 1. Consistently, all the sequence assignments in the paired houses led to losses. The assignments in the unpaired houses remained neutral. The total duration of the counterconditioning stage was equivalent to the conditioning stage (~32 minutes).

### Binary choice probe 2

This stage was identical to the previous Binary Choice Probe. It was used to confirm that counterconditioning led to a reduction in preference toward the paired houses.

### Fractals liking ratings test 2

This test was identical to the previous Fractals liking ratings and used to assess changes in liking toward the stimuli that assembled the paired houses compared to the stimuli that assembled the unpaired houses following counterconditioning.

## DAY 3

### Reinstatement

Upon their arrival to the laboratory on Day 3, participants from both groups performed a reinstatement task by re-exposure to the UCS ^29^. This was designed to reinstate the preference induced by the original Day 1 appetitive associations. Participants were instructed that they will take a short simulated tour, similar to the preceding days, except that the simulated houses will appear with no background wallpaper (i.e. without the fractals). During the task, participants entered 6 no-background houses, all yielding monetary gain, similar to the paired houses on Day 1. Consistently, all sequence assignments led to wins. The total duration of the reinstatement task was ~1.5 minutes.

### Binary choice probe 3

This stage was identical to the previous Binary choice probes. It was used to assess whether and to what extent, the reinstatement procedure recovered (i.e., increased) the preference toward the paired houses.

### Fractals liking ratings test 3

This test was identical to the previous Fractals liking ratings. It was used to assess changes in liking toward the stimuli that assembled the paired houses compared to the stimuli that assembled the unpaired houses following the reinstatement procedure.

## POST EXPERIMENT TASKS

For a detailed description of the post experiment tasks, see Supplementary Information. Briefly, we examined participants’ loss aversion propensity, using a modified version of a lottery choice task ^30,31^. Then we tested their recognition and contingency awareness of the stimuli used in the experiment. Finally, to comply with the Binary choice probes instructions, participants conducted another short simulated tour, where they entered three houses, pseudo randomly drawn from their choice trials during the Binary choice probes.

### Statistical Analysis

Binary choices between CSs that were obtained for each participant at the end of the first and the second days, and were used to determine whether to further analyze their results. Since reinstatement could be seen only after successful conditioning and counterconditioning, as manifested by preference of the paired CSs in Day 1, and its abolition in Day 2, we only included data of participants who exhibited learning of these contingencies. We set the learning criteria as follows: **1)** Successful conditioning manifested as paired choice proportion of 0.55 or higher. The rationale for this criterion was that we carefully designed an algorithm to assign the stimuli in a manner that is relatively immune to baseline preference bias and thus conceptually considered more than 0.5 choice proportion (chance level) as an indication of learning. In practice we took an interval of another 0.05 to conclude sufficient learning. In reality, the rate of almost all learners was much higher (see Results); **2)** Successful counterconditioning manifested as a lower choice proportion compared to conditioning. Here, we did not use a specific value since a subjective reduction in individual preference rates (and not an absolute value) is the most suitable indicator for this effect. Participants who failed to meet these conditions were excluded from further analyses (see Results).

We performed a logistic mixed-effects analysis between groups (Retrieval, No-Retrieval) and stage (conditioning, counterconditioning, reinstatement) with preference toward the CS+s stimuli as the dependent variable. As fixed effects, we entered group and stage (with interaction term) into the model. As a random effect, we had intercepts for participants. Specifically, we were interested in the interaction between the groups and the last two stages (counterconditioning and reinstatement). The binary choice outcome on each trial was marked as 1 or 0, according to participant’s choice of the house paired with monetary gain/loss or not. Choice trials, in which participants failed to respond within the given two seconds, were excluded from the analysis (35 observations, 0.65% of the data). On average 107.3 (SD = 1.28) observations were obtained for each participant.

To assess changes in liking ratings, we measured the difference between the average liking ratings of fractals that assembled the CSs+ compared to the average ratings of fractals that assembled CSs-. Then, using a mixed-effects linear regression analysis, we regressed this index on the variables: group (Retrieval, No-Retrieval), and stage (baseline ratings, conditioning, counterconditioning, reinstatement), with an interaction term. As a random effect we had intercepts for participants. We were particularly interested in the interaction between the groups and the last two stages (counterconditioning and reinstatement).

Recognition scores were calculated as the proportion of correctly recognizing whether a stimulus participated in the experiment or not. Contingency awareness for the winning houses was indexed as the proportion of correctly recognizing whether a stimulus that was included in the experiment was paired with monetary gain. These memory indices as well as the loss aversion index were compared between the groups using an independent samples t-test. Contingency awareness for the losing houses was not collected for all participants due to a technical problem and thus was not analyzed. Additional two participants did not perform any part of the Recognition and contingency awareness task.

We pre-determined our sample size (and pre-registered it) to be consistent with those used in previous similar studies. All regression analyses were carried out using lme4 package in R programming language (R Foundation for Statistical Computing, Vienna, Austria). P-values for the linear regression analysis were estimated using Satterthwaite approximation implemented in the lmerTest package.

Pre-registration: The procedure, hypothesis, sample size, dependent variables and analysis methods of this study were pre-registered (https://osf.io/un7y6/). To analyze the binary choice outcomes, instead of using a mixed-model ANOVA as originally planned, we performed a logistic mixed-effects analysis as described above since it is the correct analysis for this data. For the liking ratings outcomes we performed the mixed-model ANOVA-equivalent mixed-effects linear regression.

## Results

Fifty participants (25 in each group) out of 84 (41 in the Retrieval group and 43 in the No-Retrieval group) met all learning criteria and were included in the analyses. Nineteen participants did not demonstrate sufficient conditioning while 15 participants demonstrated sufficient conditioning but not counterconditioning. The overall average paired houses (CS+s) choice proportion for participants that were included in the analysis was 0.89 following conditioning and 0.37 following counterconditioning (0.34 in the Retrieval group and 0.41 in the No-Retrieval group).

### Binary choice probe (preference)

We used a mixed-effects logistic regression analysis to study the interaction between the learning stages (Stage factor) and the memory retrieval manipulation conducted prior to the counterconditioning stage (Group factor), measured by participants’ CSs choices (Table 1).

**Table 1.** Logistic regression analysis of CS+s choices as explained by Stage, Group and their interaction.

|  | **B** | **SE** | **OR** | **95% CI OR** | **p** |
| --- | --- | --- | --- | --- | --- |
| Intercept | -0.36 | 0.30 | 0.70 | [0.39, 1.25] | 0.2275 |
| Conditioning (day 1) | 3.31 | 0.16 | 27.43 | [20.19, 37.27] | 0.0000 |
| Reinstatement (day 3) | 1.96 | 0.12 | 7.09 | [5.56, 9.03] | 0.0000 |
| Retrieval Group | -0.44 | 0.43 | 0.64 | [0.28, 1.49] | 0.3032 |
| Conditioning X Retrieval Group | -0.07 | 0.21 | 0.93 | [0.61, 1.42] | 0.7455 |
| Reinstatement X Retrieval Group | -0.59 | 0.17 | 0.55 | [0.4, 0.77] | 0.0005 |

Counterconditioning (day 2) and No-Retrieval group were used as the reference categories for the Stage and Group factors, respectively.

Analysis of preference for the CSs showed a significant Group × Stage interaction (χ^2^(2) = 14.28, p = 0.0008). Following conditioning (Day 1), participants in both groups preferred the CS+s (Fig. 3), with no difference between the groups as expected, since there was no different group treatment in this phase (p=0.2542). Following counterconditioning (Day 2) participants tended to avoid the CS+s (Fig. 3), as reflected by Stage effects for conditioning and counterconditioning among both groups (Retrieval: OR = 25.59, 95% CI [19.32, 34.26], p < 2E−16; No-Retrieval: OR = 27.43, 95% CI [20.30, 37.54], p < 2E−16). Furthermore, we found no Group effect for the counterconditioning stage (p=0.3036), or a specific interaction between Group and the first two stages (conditioning and counterconditioning) (p=0.7456) as expected, since the retrieval manipulation was not hypothesized to affect counterconditioning.

Re-exposure to the appetitive UCS (Day 3) reinstated the original preference in both groups, resulting in a tendency to prefer the CS+s houses again (fig. 3). This was reflected in a Stage effect between counterconditioning and reinstatement showing greater preference towards the CS+s houses in both groups (Retrieval: OR = 3.93, 95% CI [3.15, 4.93], p < 2E−16; No-Retrieval: OR = 7.09, 95% CI [5.58, 9.06], p < 2E−16).

**Figure 3.**
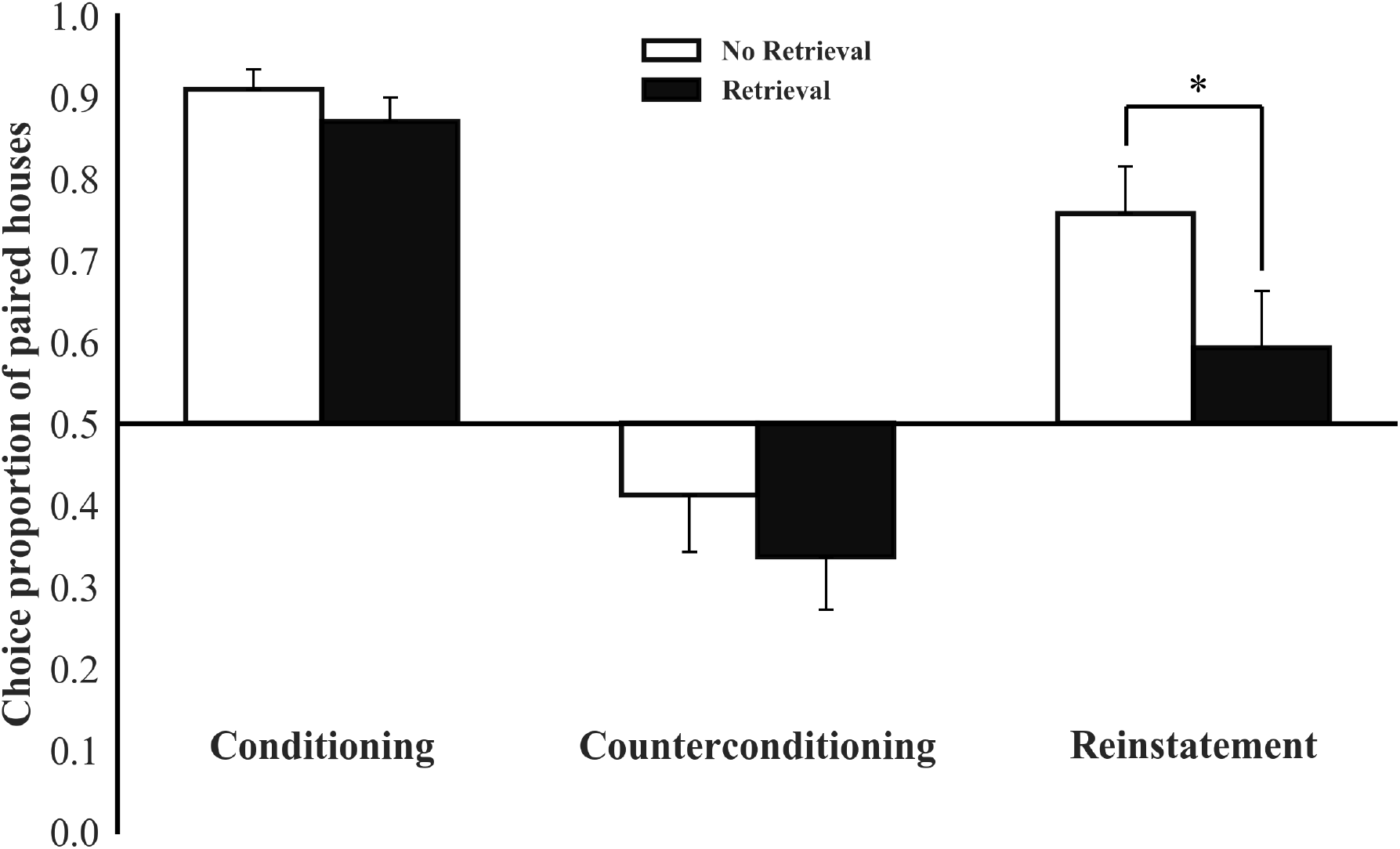
Retrieval prior to counterconditioning reduces the reinstatement of binary choice. Mean proportion of trials during which participants chose paired houses (CS+) over unpaired houses (CS-) following conditioning, counterconditioning and reinstatement. Chance level was 0.5, error bars represent standard error of the mean (SEM). *p<0.05; n=25 per group.

The most interesting effect was that in the Retrieval group, preference towards the reinforced houses was significantly less reinstated (Fig. 3), as supported by a Stage x Group interaction effect between counterconditioning and reinstatement (OR = 0.55, 95% CI [0.40, 0.77], p = 0.0005) and in a Group effect for the reinstatement stage (OR = 0.36, 95% CI [0.15, 0.83], p = 0.0163). These results suggest that the retrieval of appetitive memories before their counterconditioning reduces their capacity to influence preferences.

### Liking Ratings

We used a mixed-effects linear regression analysis to study the relationship between the learning stages (Stage factor), including the baseline, and the memory retrieval manipulation conducted prior to the counterconditioning stage (Group factor), as manifested through changes in liking toward the fractals that used to assemble the CS+s compared to the fractals that assembled CS-s (Supplemental Table S1). In contrast to enhanced preference for the paired houses, we did not find a Stage x Group interaction (χ^2^(3) = 1.18, p = 0.7575) nor a Group effect within any of the stages (p’s > 0.3617). Thus, we excluded the Group factor from the model, analyzing the Stage effect on liking across all participants. As illustrated in Supplemental Fig. S1, and consistent with the binary choice results, following conditioning (Day 1), the difference in liking ratings toward the CS+s compared to the CS-s increased significantly (β = 0.52, t(147) = 7.428, p = 8.3E-12) and reduced following counterconditioning (Day 2) (β = −0.46, t(144) = −6.638, p = 5.73E-10), approximately back to baseline. However, the reinstatement procedure (day3), did not reinstate the differences in liking observed following conditioning (β = 0.12, t(144) = 1.741, p = 0.0839), possibly suggesting that the limited delivery of the UCSs (a small quantity for a short duration) was not sufficient to reinstate the evaluation toward the stimuli that assembled the CSs. Nevertheless, it is worth mentioning that analyzing the trends within each group, without excluding the Group factor from the model, yields results that correspond with the general trend of the preferences results. This is manifested as a significant greater difference in the liking ratings towards the CS+s compared to the CS-s following reinstatement relative to the baseline (β = 0.24, t(144) = 2.403, p = 0.0175) and a marginally significant difference relative to counterconditioning (β = 0.18, t(144) = 1.845, p = 0.0671) in the No-Retrieval group, indicating a potential recovery of the liking towards the CS+s. In contrast, we found no differences between the equivalent stages within the Retrieval group (reinstatement vs. baseline: β = 0.11, t(144) = 1.153, p = 0.2510; reinstatement vs. counterconditioning: β = 0.06, t(144) = 0.601, p = 0.5490), suggesting that the original memory was not recovered following the reinstatement stage.

### Loss aversion

Two participants (one in each group) were excluded from this analysis due to choice-inconsistency, namely multiple switching points. No difference was found in loss aversion between the groups (t (46) = −0.93, p = 0.3564) with an average score of ~5.

### Recognition and winning houses contingency awareness

In the recognition task participants in both groups correctly recognized ~97% of stimuli with no difference between groups (t (46) = −1.40, p = 0.1685). Contingency awareness for winning houses in both groups was not as high (~76%) with no difference between groups (t (46) = −0.09, p = 0.9299).

## Discussion

We developed a novel multi-stage procedure to update appetitive memories in humans using aversive counterconditioning following memory retrieval. We show that induction of aversive counterconditioning shortly after retrieving an appetitive memory, reduces the reinstatement of appetitive preferences in humans. Importantly, recovery of an appetitive cue-monetary gain memory was diminished only when counterconditioning was given after memory retrieval. The evidence shown here for the ability to update human appetitive memory was obtained using a controlled laboratory procedure, targeting memory dynamics, all induced under laboratory settings in healthy participants. This extends previous attempts demonstrated on pre-existing alcohol memories of human hazardous drinkers ^15^. Furthermore, the current paradigm provides a successful translation of the retrieval-counterconditioning behavioral procedure, previously studied in animal models ^17,32^.

As opposed to the preference index, the relative liking towards the stimuli that consisted of CS+s, used as an additional index that supposedly represents the changes in evaluation of CSs, was insufficiently reinstated following the reinstatement procedure in both groups. As the effect associated with memory update could only be manifested in our procedure as a relative decrement in reinstatement, we could not assess it for the liking index. This liking index, as oppose to the preference index, was not contingent with participants’ explicit monetary prospects considerations, as the ratings were not linked with any future monetary outcome. Thus, it is possible that the intensity of the brief reinstatement was not sufficient. This could be attributed to the steadier nature of acquired evaluation, as manifested for instance in its high resistance to extinction (for review see in De Houwer et al.^33^).

A plausible explanation for the reduced reinstatement we observed is that the memory retrieval manipulation led to destabilization of the appetitive cue-monetary gain memory. This memory destabilization allowed the aversive cue-monetary loss association to incorporate into the appetitive memory trace, during memory reconsolidation. Thus, the memory was altered, consequently reducing reinstatement. Goltseker et al.^17^ recently established a similar procedure in mice, in which a cue-cocaine memory was first formed in a conditioned place preference (CPP) paradigm, and then counterconditioned with lithium chloride-conditioned place aversion (CPA) following memory retrieval. Similar to our present results, the retrieval-counterconditioning in mice prevented the reinstatement of appetitive cocaine-CPP by a cocaine prime ^17^. Critically, in the mouse study, when counterconditioning was given before, 5 hours after, or without memory retrieval, it failed to prevent reinstatement ^17^. Moreover, this effect was long-lasting ^17^. Together, our findings here alongside the mice study by Goltseker et al.^17^ suggest that the retrieval-counterconditioning procedure involves memory reconsolidation mechanisms.

Aversive counterconditioning has been suggested to be more potent than extinction in suppressing appetitive memories ^34,35^, and yet its effect may be temporary ^34,36,37^. Here, we show that when induced after a brief memory retrieval, aversive counterconditioning can suppress the reinstatement of appetitive memories, which is typically seen after counterconditioning. Interestingly, based on the memory recognition data we collected, this procedure neither affected the ability to explicitly recognize the stimuli that played a role in the task, nor the awareness to which stimuli yielded monetary gain during conditioning. We also show that the effect was not due to differences in loss aversion between the groups, which could have potentially explained the differences in reinstatement rates.

In addition to memory updating, the paradigm we constructed provides a valuable demonstration of the processes of counterconditioning and reinstatement. Previous human studies on aversive counterconditioning with laboratory-formed memories, typically employed counterconditioning immediately following conditioning ^35,38–40^, thus targeting short-term, pre-consolidated memories. Under such conditions, it is possible that aversive counterconditioning interfered with the consolidation of the appetitive memory. In contrast, in the present study, a one-day interval between conditioning and counterconditioning assured that the appetitive memory was already consolidated at the time of counterconditioning.

Reinstatement of the appetitive memories was reduced in the Retrieval group, but not completely abolished. These findings are in line with previous reports, showing that disruption of memory reconsolidation, most frequently leads to suppression of the original behavior, rather than to its complete abolition ^6,8,16,41^. It is possible that stronger aversive counterconditioning, and/or more efficient memory retrieval (e.g., by expectancy violation), could have led to a stronger suppression, or even complete abolition, of the reinstatement effect. For example, a previous human study used two consecutive days of retrieval-extinction training to decrease craving of abstinent heroin users ^14^. Das et al. demonstrated that maximizing expectancy violation (prediction error) during memory retrieval before counterconditioning led to a substantial reduction in measures of pre-existing alcohol memories of human hazardous drinkers ^15,42^.

Previous studies demonstrated that memory disruption or updating can attenuate relapse and alleviate symptoms of related disorders ^6,8,14,15,43–46^. However, others failed to find such effects with pharmacological (e.g.,: in fear conditioning – see refs ^47,48^, in drug addiction – see refs ^49,50^ or behavioral manipulations (e.g.,: see refs ^51–54^). The age and strength of the memory, as well as the manipulation used to reactivate it ^8,55^, were all pointed out as potential boundary conditions. This may complicate the translation of reconsolidation-disruption manipulations to human studies in clinical populations ^56^, and therefore our basic-science procedure allows smoother transitional process.

Maladaptive processing of associative memories has been implicated in several neuropsychiatric disorders, including addiction ^57,58^, OCD ^59^, PTSD ^60–62^ and phobias ^60^. Specifically, the reinstatement capacity of associative memories is widely responsible for the persistence and relapsing nature of these pathologies, and for the transient effects of contemporary psychotherapy. For example, relapse in addiction is strongly related to the reinstatement of drug-associated memories ^63^. Thus, a laboratory procedure to study the reinstatement phenomenon has promising clinical implications. While previous studies mainly focused on reinstatement of aversive memories (mainly fear conditioning) ^2,64^, we provide here a unique basic-science procedure that successfully induces the reinstatement of appetitive memories.

Finally, the procedure we established here is characterized by a modular structure where it is possible to selectively control and manipulate each of the stages and their different components. Exploiting this property may enable the utilization of our procedure to study similar dynamics of aversive memories or other key mechanisms in the dynamics of associative memories, compare different interventions, reinforcers and measurements, and study the underlying brain mechanisms.

In conclusion, in the past decade, several studies aimed to behaviorally interfere with the reconsolidation of associative memories. Understanding the dynamics of associative memories and their updating mechanisms holds promising clinical implications. Although there is evidence for memory alteration, the updating process is seemingly complicated, and has yet to be fully characterized. Our study, conducted under laboratory-controlled settings, successfully demonstrates appetitive memory updating in humans when employing a novel retrieval-counterconditioning behavioral procedure. Furthermore, it can be used for a wider inquiry of associative memory dynamics, relevant boundary condition and the neural mechanisms involved in the process.

## Supporting information

## Data availability and code sharing

All data and analyses codes are available on osf: https://osf.io/un7y6/

## Acknowledgement

This work was supported by the Israeli Science Foundation (ISF number 1798/15 and 2004/15) and the European Research Council (ERC) under the European Union’s Horizon 2020 research and innovation program (grant agreement n° 715016) granted to Tom Schonberg. We would like to thank Jeanette Mumford for statistics advice.

Author contribution: RG, SB and TS designed the research; RG performed the research; RG, SB and TS analyzed data and wrote the paper.

