## Supplementary material for "Counterconditioning following memory retrieval diminishes reinstatement of appetitive memories in humans"

### Supplementary information

Apparatus

Participants sat in one of two designated testing rooms and wore noise cancelling headphones. Participants in the No-Retrieval group, performed the first fifteen minutes of the second day in a third room, located in an adjacent building.

Algorithm to avoid a baseline color preference effect (implemented following Baseline fractals liking ratings o determine stimuli assignment)

For each participant, the scores of all stimuli rated were averaged. The three stimuli of each color (red, green and blue) that were the closest to the mean of all ratings were chosen to be used as the nine CSs (6 CS- and 3 CS+). Then, the average score for each CS color set was calculated. The six stimuli that belonged to the colors with the highest and lowest averages were used as CS-, and the three stimuli of the color with the intermediate average assembled the CS+. This method maximized the probability that the CS+ color is not the most liked or disliked color by each participant.

Sequence assignment (implemented during the tasks that included house visits)

Participants were instructed that sometimes, while they will be inside a house, an assignment will appear, where they will be requested to enter a sequence of letters as fast as possible (see Fig. 1C) within 1.5 seconds. They were informed that this sequence assignment may lead to “winning” or “losing" and that the faster they will press the sequence, the more money they will win or the less money they will lose. Alternatively, this sequence assignment may be “neutral”, resulting in no monetary outcome regardless of their performance. Gains and losses ranged between 0 to 0.20 NIS (~$0-$0.06 USD), determined by the relative duration until the correct sequence was entered. The sum gained/lost gradually and uniformly increased/decreased in increments/decrements of 0.02 NIS, respectively, within the 1.5 seconds. The assignment appeared in a constant timing within all houses. The timing was designed so that, in the case of the paired houses, the assignment occurred between the two spontaneous monetary rewards.

In reality, during the conditioning stage all the assignments in the paired houses (CS+) were winning.

In the counterconditioning, consistently with the spontaneous monetary loss, all the sequence assignments in the paired houses led to losses. We anticipated that due to loss aversion [1], counterconditioning will be more potent compared to conditioning. Therefore, we set the given time in the assignments (1.5 second) to be sufficient for the majority of participants to win a greater sum of money on Day 1 than they lost on Day 2. This was achieved by tests, conducted prior to the experiment, showing that it is relatively easy to enter the required sequence in less than 750 ms (half of the given time). Consequently, the sequence assignments led to unequal gains/losses between Day 1 and 2.

In both conditioning and counterconditioning stages, all the assignments in the unpaired houses (CS-) were neutral. During the Retrieval the CS+s were neutral. During reinstatement the houses appeared with no background and all of them yielded monetary gain.

<u>Features implemented to improve the realistic narrative of the simulated tour task</u>

To improve the realistic narrative of the simulated tour, a “neighborhood” screen appeared once before the task started, presenting the different houses with roads connecting between them (see Fig. 1C). Additionally, the transition between houses was indicated by the presentation of the next house to which the participant was about to enter, simulated from the outside, with a path leading to it and a door (Fig. 1C). To open the door participants were required to press a designated keyboard button. Finally, the instructions were designed to convey a story-like description, e.g., we named the neighborhood and then used its name throughout all stages of the experiment when referring to the houses.

<u>Nature video with questions task (separating Retrieval and Counterconditioning)</u>

Following the completion of the Memory retrieval task by the Retrieval group, participants watched a short nature video (~12 minutes including instructions) to provide a temporal separation between memory retrieval and subsequent counterconditioning. While watching the video, participants were asked to answer a total of 15 multiple-choice questions regarding its content. The questions were presented on the screen at a predetermined timing (every 41.3 seconds on average, SD=23.7) while the video was playing. Participants were given 10 seconds to read and answer each question. This task was fixed in duration and ensured constant short-term memory engagement.

**POST EXPERIMENT TASKS**

Loss aversion questionnaire

We asked participants to fill in a loss aversion questionnaire to test for differences between groups and if it would have affected the subjective intensity of the counterconditioning. We used a modified version of a lottery choice task [2,3]. Participants were presented with 12 sets of binary choices and were asked to indicate what they would have preferred: 1) A lottery where they could win or lose different sums of money with a chance of 50% for either. 2) A safe no-bet alternative. Later questions had higher winning sums with no change in the losing amount (i.e., a larger expected value). Loss aversion was measured by participants' switching point (ranging from 1 to 12): the higher this switching point was, the higher the participant's loss aversion tendency. Participants who did not switch to the lottery were given the score of 13.

Recognition and contingency awareness

Participants were presented with individual fractals one by one and were asked three consecutive questions: 1) Did this picture participate in the experiment? 2) Was it used as a background of a house in which you have earned money? 3) Was it used as a background of a house in which you have lost money? Participants were asked to indicate their answer and their certainty by choosing a number between 1 to 5, where 1 is certain yes, 5 is certain no and 3 stands for not sure. The nine CS-s and CS+s together with nine new fractals were presented in a randomized order and the task was self-paced. This task was designed to assess differences between groups in explicit memory indices following our procedure. The first question was used as a standard recognition measurement and the second and third were used as indicators of contingency awareness.

In the analysis, for both indices we collapsed responses of 'I am sure it was' (1) and 'I think it was' (2) and all others were considered as rejections.

Entering chosen houses

Participants performed a short task in which they entered three houses, drawn from their choices in the three Binary choice probes, in a similar manner to the Conditioning/Counterconditioning. The houses were randomly drawn from the CS- vs. CS- comparisons. Therefore, participants entered three CS-s (which resulted in no monetary outcome). This task was implemented to resolve the Binary choice probe instructions.

### Supplemental Table

**Table S1.** Linear regression analysis of mean liking ratings difference (paired – unpaired) as explained by Stage, Group and their interaction.

| | B | SE | Beta | 95% CI Beta | p |
|---|---|---|---|---|---|
| Intercept | 0.24 | 0.33 | | | 0.4617 |
| Baseline (day 1) | -0.24 | 0.42 | 0.59 | [0.10, -0.25] | 0.5776 |
| Conditioning (day 1) | 2.13 | 0.42 | 2.96 | [0.10, 0.31] | 0.0000 |
| Reinstatement (day 3) | 0.78 | 0.42 | 1.61 | [0.10, -0.01] | 0.0671 |
| Retrieval Group | 0.1 | 0.47 | 1.01 | [0.13, -0.22] | 0.8349 |
| Baseline X Retrieval Group | 0 | 0.60 | 1.17 | [0.11, -0.21] | 0.9965 |
| Conditioning X Retrieval Group | -0.33 | 0.60 | 0.84 | [0.11, -0.27] | 0.5826 |
| Reinstatement X Retrieval Group | -0.53 | 0.60 | 0.64 | [0.11, -0.31] | 0.3803 |

Counterconditioning (day 2) and No-Retrieval group were used as the reference categories for the Stage and Group factors, respectively.

### Supplemental Figure

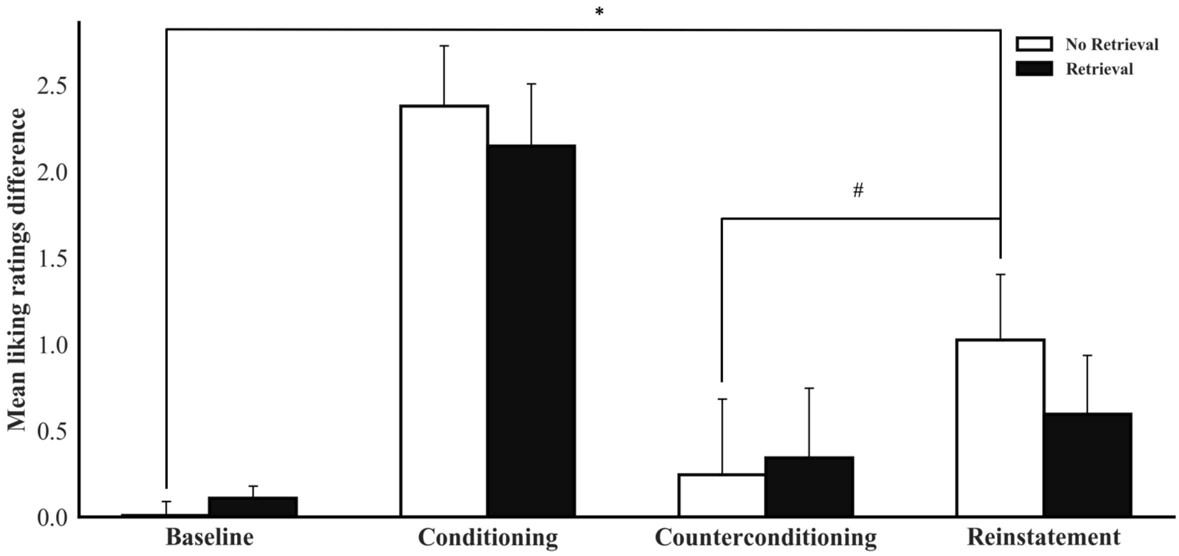


**Figure S1. Liking ratings difference.** The difference between the average ratings (on a continuous scale ranging between 0 to 10) toward the CS+s and the average ratings toward the CS-s. Error bars represent standard error of the mean (SEM). *p<0.05; #p<0.10; n=25 per group.
